# Three-Compartment Model of CAR T-cell Immunotherapy

**DOI:** 10.1101/779793

**Authors:** Brendon de Jesus Rodrigues, Luciana R. Carvalho Barros, Regina C. Almeida

**Affiliations:** Laboratório Nacional de Computação Científica; Instituto do Câncer do Estado de São Paulo

**Keywords:** Mathematical Model, CAR T lymphocytes, Memory CAR T cells, Long-term Immunity, Tumor Induced Immunosupression

## Abstract

Immunotherapy has gained great momentum with chimeric antigen receptor T-cell (CAR T) therapy, in which patient’s T lymphocytes are genetically manipulated to recognize tumor-specific antigens to increase tumor elimination efficiency. Improved CAR T cell immunotherapy requires a better understanding of the interplay between CAR T cell doses and tumor burden, administration protocol, toxicity, resistance to immunotherapy, among other features. We developed a three-compartment mathematical model to describe tumor response to CAR T cell immunotherapy in immunodeficient mouse models. It encompasses interactions between tumor cells, effector and long-term memory CAR T cells such as tumor induced immunosuppression effects, conversion of memory T cells into effector T cells in the presence of tumor cells, and individual specificities considered as uncertainties in the parameters of the model. The model was able to represent two different immunotherapy scenarios with different CAR receptors and tumor targets reported in the literature. Further *in silico* studies considering different dosing quantities and tumor burden showed that the proposed model can represent the three possible therapy outcomes: tumor elimination, equilibrium, and escape. We found that therapy effectiveness may also depend on small variations in the parameter values, regarded as intrinsic individual specificities, as T cell proliferation capacity, as well as immunosuppressive tumor microenvironment factors. These issues may significantly reduce the chance of tumor elimination. In this way, the developed model provides potential use for assessing different CAR T cell protocols and associated efficacy without further *in vivo* experiments.

## Introduction

Adoptive cell therapies have been considered a major advance in the fight against cancer, especially those associated with the hematopoietic system [27]. CAR T cell immunotherapy is an adoptive cellular therapy in which T lymphocytes are taken from the blood of a patient, genetically modified to recognize antigens expressed by the patient’s tumor, submitted to *in vitro* expansion and reinjected into the patient. Insertion of the CAR gene into T lymphocytes bestows the ability to recognize and directly attack tumor cells regardless of human leukocyte antigen presentation [32]. In 2017, the FDA approved the commercialization of two therapies with CAR T cells for the treatment of CD19^+^ B cells malignancies [16, 26]. Other target proteins have been studied recently, such as CD123 that is expressed in many hematological malignancies, including acute myeloid leukemia, Hodgkin’s lymphoma, acute lymphoblastic leukemia, among others, which makes it a potential antineoplastic target [6]. Therapy with CAR T cells has been successful in eliminating or relieving endurable types of lymphomas and leukemia. Recent experimental studies have investigated the relationship between immunotherapy with CAR T cells and the development of immunological memory in cancer [30, 5]. Using immunodeficient mouse models, [30] showed that CAR T 123 therapy can eliminate Hodgkin’s lymphoma and provide long-term immunity against recurrence of the same tumor. Immuno checkpoint blockade associated with CAR T cell therapy is also under investigation in models where CAR T cell therapy fail. Also using immunodeficient mouse models, [29] showed that tumor expressing indoleamine 2,3-dioxygenase (IDO) activity, an intracellular enzyme that has an inhibitory activity on T cells, can be better controlled by combining the CAR therapy with 1-methyl-tryptophan (1-MT), an IDO inhibitor. By the end of 2016, four different immune checkpoint blockade drugs were also approved for the treatment of lymphoma, melanoma, among other cancers. Current and future advances in the engineering of CAR and new immunologic checkpoint inhibitor drugs offer promising perspectives in the treatment of cancer [16].

Although the success of this type of therapy against hematologic cancers is promising, the mechanisms associated with failures have been reported and are subjects of recent investigations [19]. Notably, many challenges remain to be addressed to improve response rates such as minimum effective T cell dose, subtypes of CAR T cells selection, adverse effects, combination with other types of therapy, suppressive microenvironment, and patient specificity, among others. Mathematical models may contribute to the understanding of these mechanisms [5, 2, 22, 23], confronting hypotheses and testing different settings [20, 39, 24]. To this end, here we focus on the development of a mathematical model to describe the treatment of hematologic malignancy in immunodeficient mouse models with a CAR T immunotherapy and the formation of the immune memory pool. We calibrated the model parameters with *in vitro* and *in vivo* data presented in [30] for CAR T 123 therapy against Hodgkin’s lymphoma. We then used the model to retrieve a different immunotherapeutic CAR scenario, using data from [29] for CAR T 19 therapy on Raji tumors. We complemented the *in silico* study investigating different therapy outcomes depending on the relationship between the tumor burden and CAR T cell number; the therapy effectiveness due to inhibition by immunosuppressive tumor microenvironments, and intrinsic individual specificities represented by the uncertainties in the parameters of the model. Our model simulations provide insights on critical mechanisms of CAR T-cell therapy, and show potential use for assessing different CAR T cell protocols and associated efficacy, complementing and potentially avoiding further *in vivo* experiments.

## Materials and methods

### *In vitro* and *in vivo* data

We used *in vitro* and *in vivo* data published in [30] and [29]. The cytotoxic activity of CAR T cells was evaluated in a standard *in vitro* 4-hour chromium-51 release assay [13]. Bioluminescence imaging (BLI) data were used to track tumor growth time-course. We considered one BLI unit as one cell. Although we did not find any correspondence in the literature to convert BLI to cell number, BLI and total tumor cell number is directly correlated with the total number of cells as shown in [1]. All used data were extracted from those sources using the free software [11] and, for completitude, are presented in the Supplementary Material.

### Model development

In this work, we focus on the development of a three-compartment mathematical model, using ordinary differential equations, to describe the interactions between populations of a tumor, effector CAR T and memory CAR T cells. As we are dealing with immunodeficient mice, we consider that the effector CAR T cells come only from the immunotherapy, represented by populations of (activated) CAR T lymphocytes that we denote by *C*_*T*_. The population of memory CAR T lymphocytes is denoted by *C*_*M*_ and the tumor population is denoted by *T*. The change of each cell population in time depends on the balance among all factors contributing to its increase and decrease. The interaction mechanisms between model compartments are schematically described in Figure 1. Notably, we chose to individually model each mechanism in order to better characterize it.

**Fig. 1.**
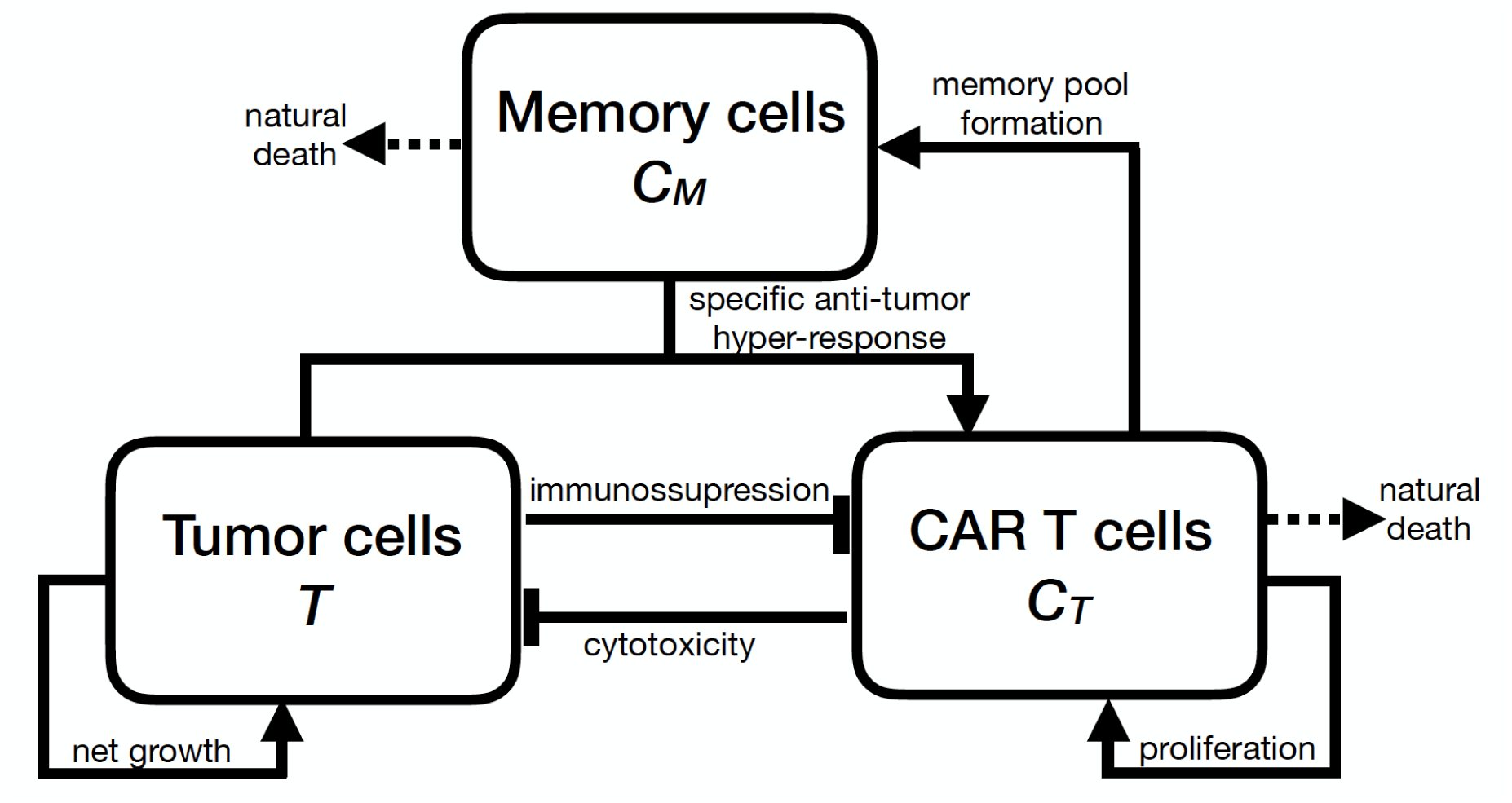
Schematic description of the model structure. CAR T cells proliferate, have a cytotoxic effect on tumor cells, differentiate into memory cells, and die naturally or due to immunosuppressive mechanisms. The long-lived memory T cells also die naturally and are readily responsive to the tumor associated antigen so that, when they interact with tumor cells, they differentiate into effector T cells, producing a rapid response of the immune system.

We assumed that a given dose of activated CAR T cells is introduced into the system as an adoptive therapy. The effector CAR T cells have antitumor activity owed to a specific chimeric antigen receptor (CAR) against an antigen expressed by the tumor [33, 38]. The overall net population balance of the effector CAR T cells depends on their spontaneous death and proliferation, inhibition by the tumor cells, and differentiation to a memory CAR T cell phenotype [34, 37]. This means that a portion of CAR T cells persists as memory T cells that keep antitumor antigen specificity and patient characteristic [21, 31]. They have a lower activation threshold, which eases the secondary response to a future tumor recurrence [35]. They also have a long half-life, providing long-term protection. In the absence of immunosurveillance, the tumor grows exponentially but we also assumed that the growth rate decreases linearly with the population size, approaching zero as the tumor population approaches the maximal tumor burden (carrying capacity) [8, 28]. Upon contact, effector CAR T cells kill tumor cells. As the rate of CAR T cell killing capacity could not be directly measured *in vivo*, we assumed the constant rate as a range of tumor cell killing by effector CAR T cells over time [14, 17, 3].

The change of each cell population in time depends on the balance among all factors contributing to its increase and decrease. Specifically, the dynamics of the effector CAR T cells is described by:

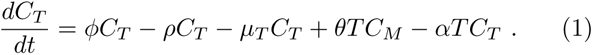

The right-hand side term specifies that effector CAR T cells undergo expansion due to proliferation at a rate of *φ*, and have half-life 1*/µ_T_*. According to the linear progression model described in [5, 12], they differentiate at a rate of *ρ* into long-term CAR T memory cells, which are assumed to provide long-lasting protection to the specific tumor/antigen. At any future time in which memory CAR T cells come into contact with the same tumor cells, they can rapidly be converted into effectors CAR T cells, readily activated to prevent tumor progression. This feature is modeled by the term *θTC_M_* represents. Finally, the term *αTC_T_* models tumor-modulated CAR inhibition that can encompasses a variety of immunosuppressive mechanisms.

The dynamics of the immunological memory, a key dynamic of the adaptive immune system [5, 36] is modeled by

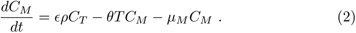

Since we are dealing with immunodeficient mice, memory T cells are formed exclusively from the differentiation of CAR T cells at a rate of *ερ*. When in future contact with the same antigen-bearing cancer cells, they immediately return to the effector CAR T cell phenotype at a per capita rate proportional to the tumor burden. In general, memory T cells have longevity, and therefore have a much lower mortality rate than the effector T cells [36], i.e., *µ_M_* << *µ_T_*.

The response of tumor cells to the CAR T immunotherapy is modeled by

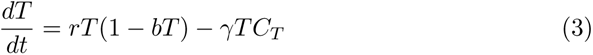

The first term in the right-hand side of (3) models the density-dependent growth of cancer cells due to the limitation of available resources in the tumor microenvironment. Tumor growth is described using a logistic function in which *r* is the maximum growth rate and 1*/b* is the carrying capacity, i.e., the maximum cell density that the available resources are capable of sustaining. CAR T cell immunotherapy acts by inhibiting tumor growth by cytotoxic action, causing a per-capita mortality rate of *γC_T_* that depends on the number of effector T cells.

The meaning of the parameters (strictly positive real numbers) is given in Table 1. Notice that model (1)-(3) has ten parameters. Their estimation ultimately defines the desired immunotherapy scenario. Of note, the linear same order terms in equation (1) can be combined to reduce the number of parameters. This procedure is particularly useful for cases where experimental data is scarce, for which a single rate may be alternatively used to reproduce the balanced effect of proliferation, death, and differentiation of effector CAR T cells. Details about parameter inference are presented in the Supplementary Material. The next section shows that the overall approach provides a framework for investigating the roles of CAR T cell dose, immune memory, immunosuppressive tumor microenvironment and individual uncertainties on the therapy response.

**Table 1.** Summary of the model parameters.

| Parameter | Meaning | Unit |
| --- | --- | --- |
| $\phi$ | $C_T$ proliferation rate | $day^{-1}$ |
| $\rho$ | Differentiation rate of $C_T$ into $C_M$ | $day^{-1}$ |
| $\mu_T$ | $C_T$ death rate | $day^{-1}$ |
| $\theta$ | Conversion coefficient of $C_M$ into $C_T$ due to interaction with $T$ | $(cell \cdot day)^{-1}$ |
| $\alpha$ | $C_T$ inhibition coefficient due to interation with $T$ | $(cell \cdot day)^{-1}$ |
| $\epsilon$ | Numerical response of the conversion of $C_T$ into $C_M$ | - |
| $\mu_M$ | $C_M$ death rate | $day^{-1}$ |
| $r$ | Maximum growth rate of $T$ | $day^{-1}$ |
| $b$ | Inverse of the tumor carrying capacity | $cell^{-1}$ |
| $\gamma$ | Death coefficient induced by $C_T$ | $(cell \cdot day)^{-1}$ |

## Results: *In silico* experiments

Mathematical equations (1)-(3) were solved numerically using the explicit fourth-order Runge-Kutta method [9]. Simulations represent CAR T cell immunotherapy performed in immunodeficient mice previously injected with tumor cells. This amounts to set the initial condition for the tumor population, *T* (0), as the injected tumor cells, and *C*_*T*_ (0) = *C*_*M*_ (0) = 0 cells. At the time the immunotherapy is given, when tumor burden has already undergone significant growth, *C*_*T*_ receives the CAR T cells dose. Cell populations are followed up to investigate tumor response and immunological memory formation.

CAR T 123 therapy eliminates HDLM-2 tumor, providing long-term protection; the immunotherapy with CAR T 19 on Raji tumor slows down its growth

We first simulate the scenario presented in [30] that consists of CAR T 123 therapy against HDLM-2 cells. Ruella et al. [30] reported that 2 × 10^6^ cells of Hodgkin lymphoma (HDLM-2) were injected into NSG mice. After 42 days, mice received *C*_*T*_ = 2 × 10 cells of CAR T 123 immunotherapy, which eliminated tumors very rapidly. Model parameters are displayed in Table S6 of the Supplementary Material. Simulation begins with *T* (0) = 2 × 10^6^ HDLM-2 cells, and tumor progresses in time until it reaches about 2 × 10^7^ cells in *t* = 42 days (Figure 2(a)). At this time, immunotherapy with CAR T 123 is performed, so that *C*_*T*_ = 2 × 10^6^ cells at *t* = 42 days. Immunotherapy rapidly eliminates the population of tumor cells in a few days, retrieving the experimental remission results presented in [30]. Our simulation also provides the dynamics of effector and memory T cells. Figure 2(a) shows that, as the population of *C*_*T*_ cells decrease, phenotypic differentiation occurs giving rise to memory T cells *C*_*M*_. Our simulation shows that effector CAR T cell populations remain undetectable until *t* = 250 days, which agrees on results presented in [30] around ten months after the injected dose. Moreover, our model indicates the presence of long-term memory CAR T cells, which slightly decline in time due to small mortality rate of *µ_M_*.

**Fig. 2.**
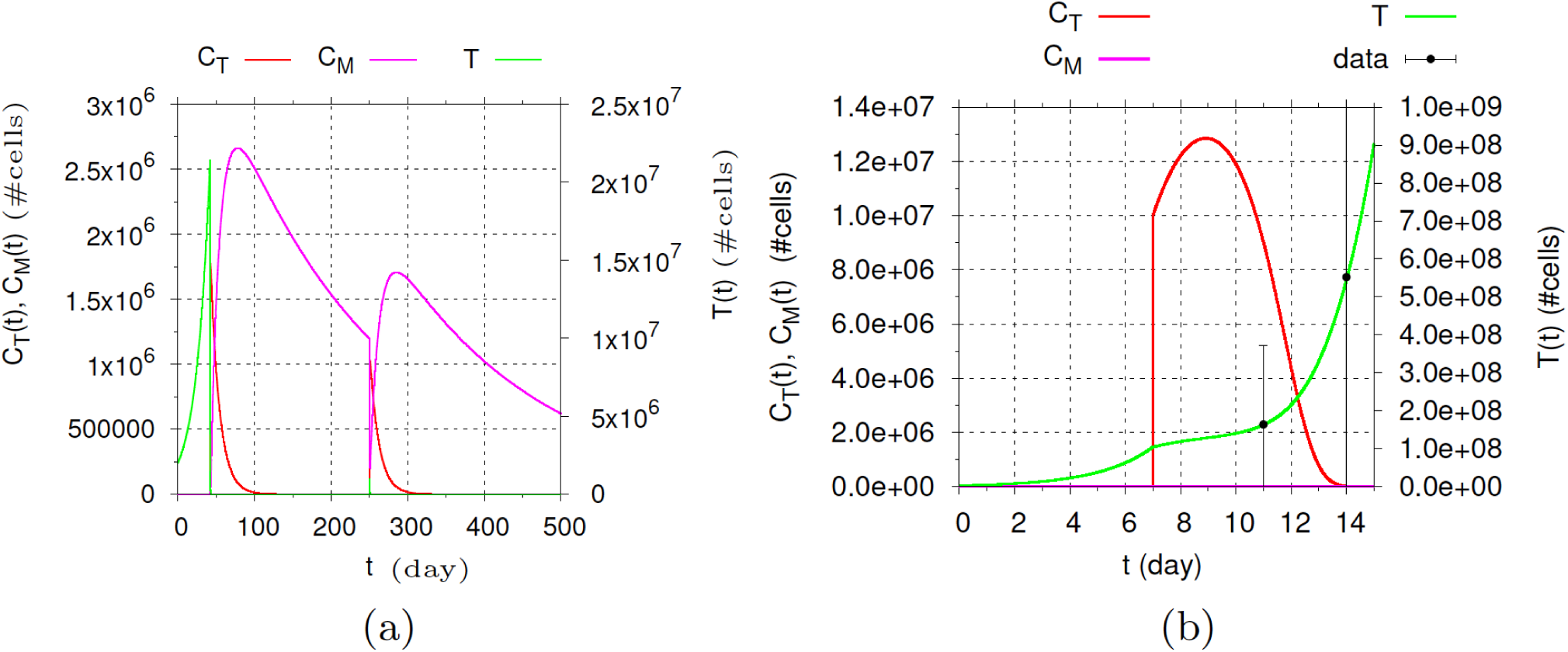
Dynamics of *T*, *C*_*T*_ and *C*_*M*_ cell populations. (a) The immunotherapy with CAR T 123 on HDLM-2 and challenge are performed at *t* = 42 and *t* = 250 days, respectively. Tumor is rapidly eliminated after *C*_*T*_ is introduced in the system. Soon after there is a decay of *C*_*T*_, which is partially converted into *C*_*M*_. Tumor remains undetectable until day 250. Simulation is continued by carrying out a challenge at day 250. Upon contact with new tumor cells, *C*_*M*_ is converted into *C*_*T*_, which rapidly eliminates the tumor. Afterwards, immunological memory is partially recovered. (b) The immunotherapy with CAR T 19 on Raji-control is performed at day 7. There is an expansion of effector T cells, which can reduce growth but not eliminate the tumor. Effector T cells are practically extinct at the end of the simulation. There is no formation of memory CAR T cells. Data extracted from [29].

In an additional experiment, Ruella et al. [30] demonstrated the formation of the immune memory by challenging previously treated mice with 1 × 10^6^ HDLM-2 cells at *t* = 250 days. The tumor remained undetectable, being rejected due to the re-expansion of the effector CAR T cells. To interrogate the model on this behavior, we continue the previous simulation by introducing 1 × 10^6^ tumor cells at *t* = 250 days. Figure 2(a) shows how the model answers such a challenge. The presence of tumor cells drives the conversion of *C*_*M*_ into *C*_*T*_ cells which are rapidly able to eliminate the new tumor. Afterward, *C*_*T*_ undergoes rapid decay while part of the memory T cells population is recovered. Tumor clearance remains until the end of simulation on day 500. As explained in [30], tumor rejection occurs due to the re-expansion of previously undetectable effector CAR T cells.

We next fit the model to the scenario described in [29], regarding Raji tumor and immunotherapy with CAR T 19 cells. Raji tumors are much more aggressive than HDLM tumors and express the CD19 antigen. Ninomiya et al. [29] reported that 3 × 10^6^ Raji tumor cells were injected in SCID/Beige mice and therapy with 1 × 10^7^ cells of CAR T 19 cells was given in day 7, which did not eliminate the tumor but could control its growth. This scenario is simulated with the estimated parameters displayed in Table S6 of the Supplementary Material. Beginning with *T* (0) = 3×10^6^ cells, the tumor reaches almost 1 × 10^8^ cells at day 7 when *C*_*T*_ = 1 × 10^7^ cells of CAR T 19 cells is introduced. Retrivieng the results presented in [29], the immunotherapy is able to reduce the tumor growth rate but not eliminate it, and tumor cell population reaches 6 × 10^8^ cells at day 14, as shown in Figure 2(b). The effector T cells undergo an expansion of about 30% on day 9, from which they decrease to extinction, representing the T cell time course reported in [29]. The immunotherapy dose is not enough to lead to the formation of memory CAR T cells.

### Insights on CAR T 123 dosing strategy into the elimination of HDLM-2 tumors

The model was used now to investigate how the relationship between tumor burden and CAR T cell dose and injection protocol impact therapy outcomes. To first assess how the dose interferes with the response to the CAR T 123 immunotherapy, we perform three different simulations with therapeutic doses of 1 × 10^6^, 5 × 10^5^ and 2 × 10^5^ *cells*. We use the same scenario described in Figure 2(a) and same model parameters, keeping the initial tumor burden equal to *T* (0) = 2×10^6^ cells. The resulting dynamics are shown in Figure 3(a)-(c). A CAR T dose of 1 × 10^6^ *cells* can perform tumor elimination, although the level of memory T cells at *t* = 200 *days* is smaller than that in the case presented in Figure 2(a). Higher CAR T cell dose generates greater immunological memory T cell pool. On the other hand, reducing the dose of CAR T cell to 5 × 10^5^ *cells*, the tumor is not completely eliminated. It undergoes an intense decrease but resumes growth at day 150, eventually reaching a state in which it does not grow or shrink significantly, wherein the tumor is reduced to a very small (but not zero) value as depicted in Figure 3(b). In this equilibrium state, both *C*_*T*_ and *C*_*M*_ s are non-zero, and therefore there is the coexistence of the three cell populations. This is a typical configuration of tumor dormancy. Finally, reducing, even more, the CAR T dose to 2×10^5^ *cells*, the tumor escapes from the immunotherapy. Figure 3(c) shows that the tumor is initially reduced by therapy (not visualized because of the scale) but resumes growth and reaches the carrying capacity at around day 300. There is a complete and rapid extinction of the CAR T population and no formation of memory CAR T cells.

**Fig. 3.**
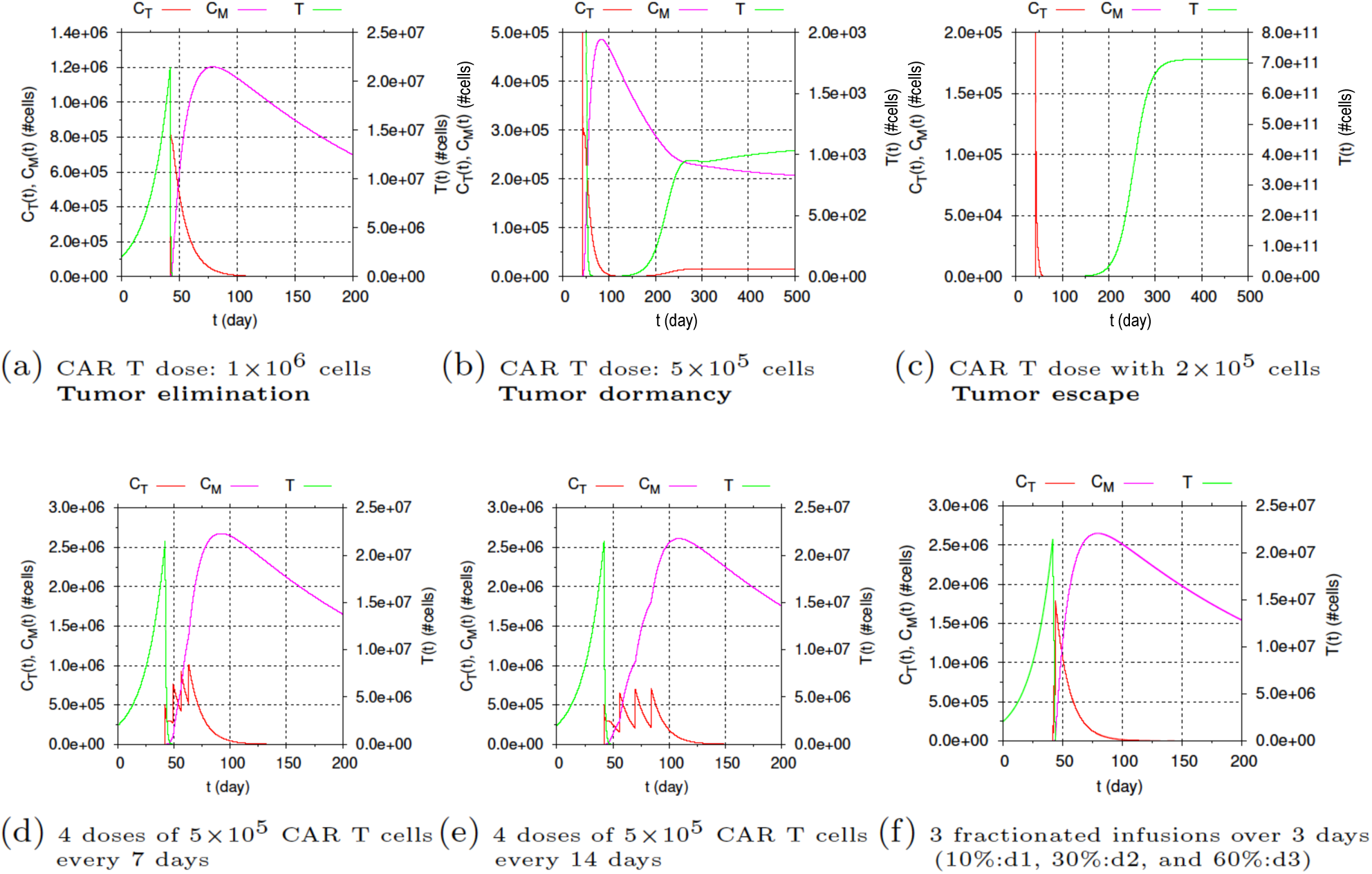
*In silico* prediction of the immunotherapy response to different CAR T cell dose. Initial HDLM-2 tumor burden amounts 2 × 10^6^ *cells*. Top row: (a) with 1 × 10^6^ CAR T cells injection, tumor elimination occurs around day 100; around 7 × 10^5^ memory cells remain at *t* = 200 *days*; (b) half of the previous CAR T cell dose (5 × 10^5^) induces a strong decline in the tumor burden, together with a decrease in the number of CAR T cells. However, tumor rapidly resumes growth. After day 250, the three cell populations slowly change towards an equilibrium state, in which a small pool of tumor cells coexists with the CAR T and memory T cell populations; (c) 2 × 10^5^ CAR T cells dose is not able to control the tumor, which escapes and reaches the carrying capacity at day 350. The fast decay of CAR T cells prevents the formation of a memory cell population. Bottom row: the total CAR T dose of 2 × 10^6^ *cells* is fractionated into four equal portions and administered every (d) 7 days or (e) 14 days; (f) the dose is fractionated into 3 infusions of increasing dose values over 3 days. In all cases (d)-(f), the tumor is eliminated in a few days, followed by a decrease of the effector T cells. Fractionated infusions lead to the formation of memory T cells, although the quantity depends on the rest time between doses.

Although not shown, it is worth remarking that those three tumor responses of elimination, equilibrium, and escape can be reached by fixing the CAR T dose and increasing the tumor burden.

The next experiment explores the alternative possibility of CAR T cell dose fractionation. We select the same scenario described in Figure 2(a) with 1-time infusion of 2 × 10^6^ *cells*, which promotes tumor elimination. Firstly, simulations are performed dividing the total dose into four equal fractions, infused at every seven or fourteen days. Figures 3(d) and 3(e) show that the dosing split does not interfere on the tumor elimination, which occurs in few days. Of note, a single dose of 5×10^5^ CAR T cells is not able to eliminate the tumor burden, as shown in Figure 3(b). While in a single infusion case tumor decreases but resumes growth until reaching equilibrium, the used fractionated infusions prevent tumor regrowth. As well as in Figure 2(a), immunological memory is formed, and the peak of memory cells is similar to that of single total dose infusion, although a certain delay is observed due to dose fractionation. Such delay ultimately yields a greater formation of immunological memory on day 200. Specifically, the number of memory CAR T cells at that time is 10% and 17% for 7 and 14 days rest time between doses, respectively. Although this feature could be seen as an advantage towards fractionated infusions, long rest periods between doses cannot be used because CAR T cells do not survive in culture medium for such long times. Alternatively, a simulation is performed for a more realistic fractionated immunotherapy, as investigated in [10]. In that work, patients with relapsed or refractory CD19+ acute lymphoblastic leukemia were treated with three fractionated infusions over 3 days with increasing doses (10%:d1, 30%:d2, and 60%:d3). It was shown that such treatment protocol does not compromise effectiveness while reducing toxicity effects. Figure 3(f) shows the *in silico* predictions using this protocol. Like in a 1-time infusion protocol shown in Figure 2(a), tumor is rapidly eliminated, effector T cells vanish in 100 days while immunologic memory amounts for 1.5 × 10^6^ cells at day 200.

### How do parameter uncertainties impact the elimination of HDLM-2 tumors?

We now use *in silico* experiments to investigate how parameter uncertainties impact the CAR T 123 immunotherapy outcomes. We selected the scenario depicted in Figure 2(a): HDLM-2 tumor burden of *T* (0) = 2 × 10^6^ cells, CAR T 123 dose of 2 × 10^6^ cells introduced at day 42, and the set of parameters depicted in Table S6 of the Supplementary Material. Model simulation under these conditions leads to tumor elimination. We then considered that parameter values are uncertain. Recalling that parameter values were estimated for the scenario defined in [30], uncertainty can be regarded as associated with individual characteristics of each mouse, tumor cell lineage, and CAR T cell donor. Since we did not have any evidence of such uncertainties, we assumed that each parameter is a random variable with uniform distribution in the range limited by 20% of the reference values indicated in Table S6. We built 10,000 sets of parameters from randomly selecting samples of the parameter distributions. We then simulated the model for each set of parameters in order to assess if small random variations in the parameter values can impact tumor elimination. We verified that they indeed impact the therapy outcome, and we registered the elimination, equilibrium or escape responses. The simulations indicated that the uncertainties in the parameters can drastically reduce the chance of therapy success: of the 10,000 cases considered, the therapy was successful in only 5% of them (507 cases). The equilibrium and escape responses occurred in 18% and 77% of the total 10,000 cases, respectively. Overall, the performed simulations indicate that tumor response is very sensitive to parameter uncertainties, and tumor escape is most likely to occur. To assess how each parameter value impacts these responses, we then built the three heatmaps shown in Figure 4, that displays the frequency of occurrence of each outcome (elimination, equilibrium, and escape) with respect to the parameter values. The inspection of these plots indicates that the elimination heatmap is more heterogeneous while the escape one is more homogeneous. The latter indicates that all parameters contribute more uniformly to this outcome, meaning that none of the parameters plays a significant differential role in tumor escape. On the other hand, the more heterogeneous occurrence frequencies in the elimination heatmap allow identification that the tumor proliferation (*r*), CAR T cell inhibition (*α*), and CAR T cell proliferation (*φ*) and death (*µ_T_*) are the most influential for tumor elimination.

**Fig. 4.**
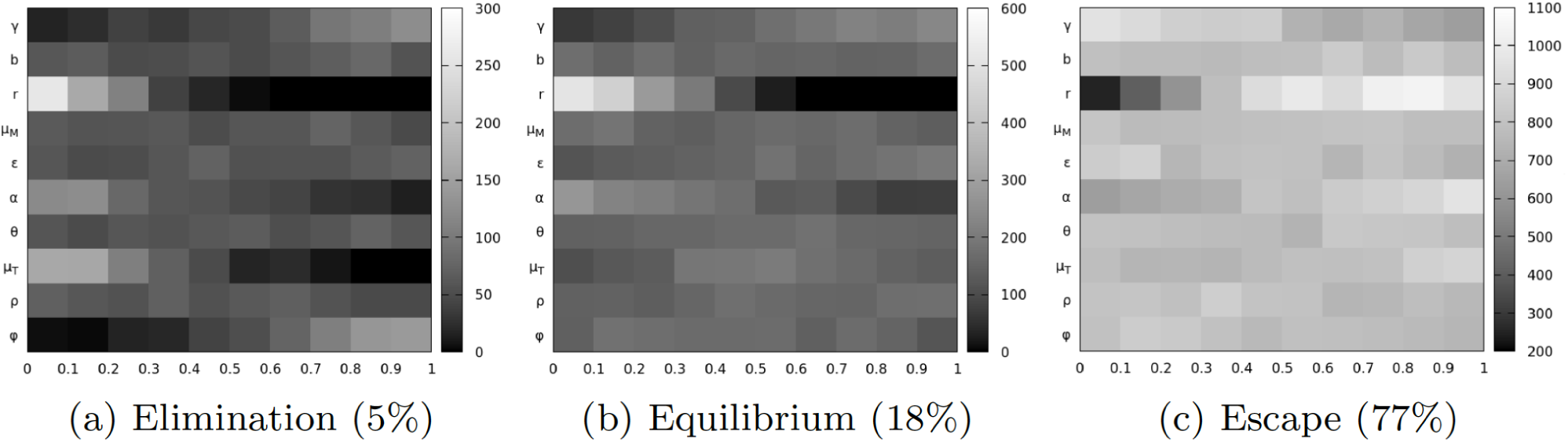
Frequency of occurrence of elimination (a), equilibrium (b), and escape (c) of the tumor for the scenario in which the initial HDLM-2 tumor burden is 2 × 10^6^ *cells* and the immunotherapy with 2 × 10^6^ *cells* CAR T is performed at day 42. Darker colors indicate lower frequency. All parameters are assumed to be uncertain, being uniform distributed random variables with range limited by 20% of the reference values indicated in the Supplementary Material, **Table S6**. Horizontal axes are associated with the normalized parameter values. We evaluated 10,000 cases by randomly sampling the parameter space. Tumor escape is more likely to occur, followed by the equilibrium and elimination. The respective percentage of these responses are indicated in parentheses.

### The effect of inhibitors of immunosuppressive tumor microenvironments

Our model includes the term *αTC_T_* in Equation (1) to describe tumor-modulated immunosuppressive mechanisms. Higher *α* value implies in a stronger immuno-suppressive mechanism. To check how this term allows investigating the blocking action of these mechanisms and, at the same time, how the model deals with different tumors and CAR T cells, we selected data from [29] that presents the action of CAR T 19 cell immunotherapy against CD19 + lymphoma that expresses the IDO enzyme in mice. We then considered the Raji-IDO tumor when treated with CAR T 19 cells alone or combined with 1-MT, an IDO inhibitor. We estimated *α* for these scenarios, keeping all other parameters fixed with values shown in Table S6 for the Raji tumor (control). It should be noted that, according to Figure 2C of [29], Raji-control and Raji-IDO tumor sizes on the day of immunotherapy administration are indistinguishable so that the same tumor proliferation rate was used. Figure 5 shows system responses for the Raji-IDO + CAR T 19 and Raji-IDO + CAR T 19 + 1-MT scenarios, together with the most likely values (MLEs) obtained for *α*. The smaller *α* value obtained when 1-MT was used allowed a greater expansion of CAR T cells after infusion which in turn provided a stronger control on the tumor growth than that promoted by the CAR T cells without 1-MT. Of note, in both cases the CAR T 19 dose was not able to eliminate the tumor, which eventually escapes, and there is no formation of memory CAR T cells. We can also notice the similarity of *α* values for the Raji-control + CAR T 19 and Raji-IDO + CAR T 19 + 1-MT, reflecting the ability of the 1-MT to block the immunosuppressive effect of the IDO. Thus, the model could capture the effect of the IDO inhibitor through the *α* parameter that can actually modulate the immunosuppression mechanism used by Raji-IDO tumors. These simulations show the ability of *α* in modulating immunosuppressive mechanisms displaying the potential use of our mathematical model as an adjuvant *in silico* platform to test immune checkpoint inhibitors.

**Fig. 5.**
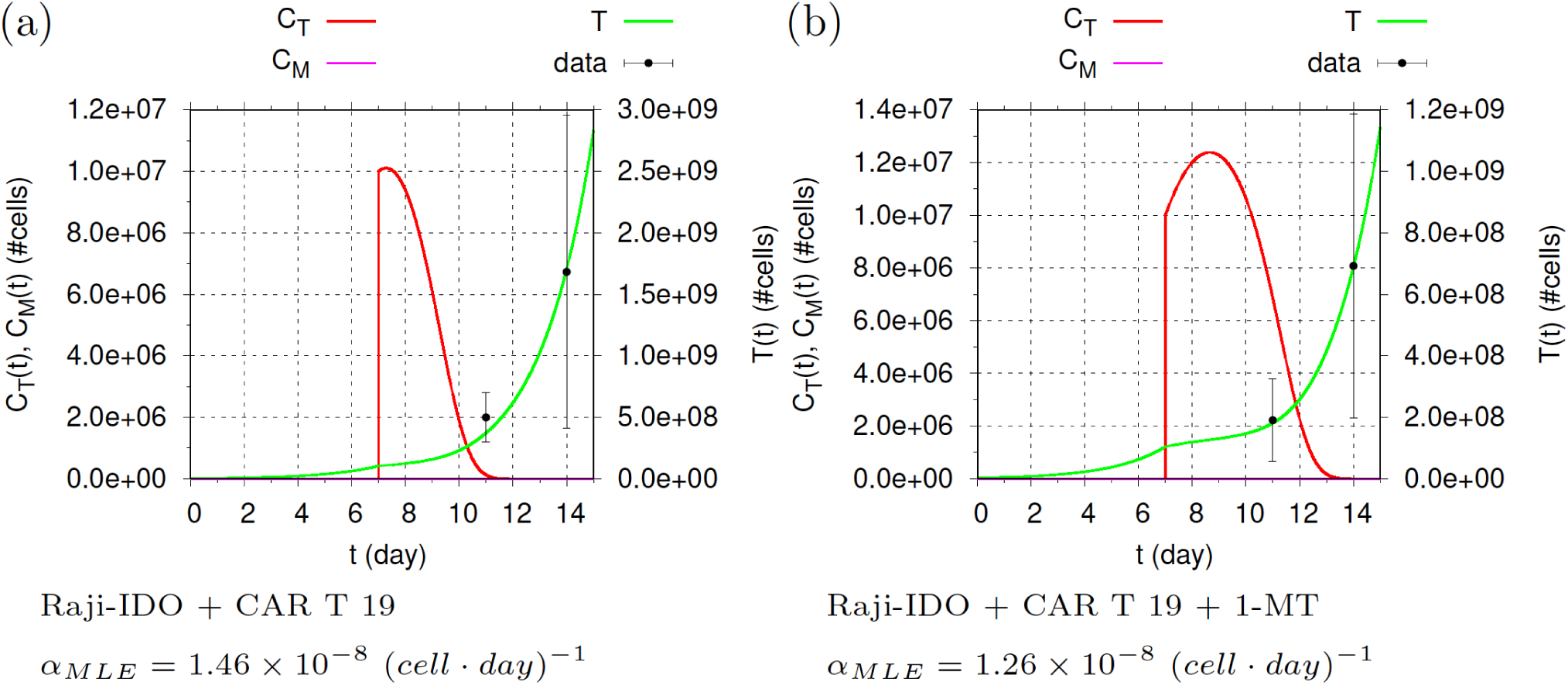
System response to: (a) CAR T 19 immunotherapy; (b) CAR T 19 immunotherapy with IDO inhibitor (1-MT). Initial Raji-IDO tumor burden is 3 × 10^6^ *cells*. At day 7, 1 × 10^7^ CAR T 19 cells were introduced and were able to reduce the velocity of the tumor growth at some extent. Such reduction was more significant when CAR T 19 therapy was combined with the IDO inhibitor (1-MT): the number of tumor cells were less than half of that without 1-MT at day 15. Model parameter *α* was estimated for these two cases, and their corresponding MLE values are indicated above. The parameter *α* was responsible to capture the effect of IDO inhibition and the 1-MT. Its value decreased for the Raji-IDO + CAR T 19 + 1-MT case, being small enough to promote a higher expansion of the CAR T cells, and ultimately leading to a more effective control on the tumor growth. However, both therapies were not able either to eliminate the tumor or build memory cells. Data extracted from [29].

## Discussion

CAR T cell therapies are spreading across hematological cancers and are already a product of big pharma companies [25]. On the road, there are new CAR designs, including new antigen targets, different CAR affinity [15], and expansion protocols [7].

We developed a three-compartment mathematical model to describe tumor response to CAR T cell immunotherapy in immunodeficient mouse models (NSG and SCID/beige) based on two published articles from literature. In a general CAR T cell therapy model, independently of the recognized antigen, we modeled different receptors as CART 19BBz and CART 123, and also different tumor targets as HDLM-2 and Raji. The HDLM-2 tumor model was used as a low proliferation, less aggressive tumor model where CAR T cell therapy is effective on tumor elimination. On the other hand, the Raji model was chosen from its high proliferation and escape from CAR T cell therapy. On the Raji model we also included explicitly immune checkpoint inhibitor molecule as IDO in order to estimate this component on CAR T/tumor cell interaction. Therefore, the model could adapt itself to different treatment and tumor scenarios. The adopted structure of our mathematical model allows identifying each individual mechanism in a more transparent way. Donor/tumormicroenvironment specificities were considered as uncertainties in the parameters of the model, which were shown to greatly impact the therapy outcome. The model was able to represent tumor elimination after immunotherapy with CAR T 123 cells even in case of a new tumor challenge due to memory T cells’ long-term protection for HDLM-2 target. The change of CAR T cells from effector to memory cells and their long-term persistence as CAR T memory cells were also demonstrated by our previous work with RS4;11 B-ALL model using 19BBz CAR T [4]. For the CAR T 19 therapy and Raji target scenarios, the model represented well the tumor dynamics with or without IDO inhibitor. We performed a few *in silico* studies to highlight how the model can be used as an adjuvant platform to contribute to a better understanding of the underlying processes. We found that the determination of the dose of CAR T for a given tumor burden is a critical factor for the success of the immunotherapy. A previous model already considered CAR T cell proliferation in response to antigen burden, but memory CAR T was not considered, neither the effect of tumor inhibition of CAR T cells [37]. Another interesting mathematical model was made upon tisagenlecleucel-treated patient data [34]. This model was adapted from a previous empirical model of an immune response to bacterial/viral infections. They captured T cell expansion, contraction, and persistence like our model does, including CAR T memory population. Their model was calibrated on patients’ data, and different from ours, no difference in dose-response was detected. They attributed this result to CAR T cell proliferation capacity *in vivo*. We partially agree, but there is a possibility that obtained data from humans do not present very different CAR T cell dose (especially including only tisagenlecleucel clinical trials). Considering mouse model data, where CAR T cell dose varies by thousands, we do observe a dose effect, especially on aggressive, high proliferative tumors as NALM-6 cell line [7, 4].

Another advantage of our mathematical model is the calculation of therapy effectiveness. Overall therapy effectiveness may depend on intrinsic individual specificities, regarded here as small variations in the model parameters’ values. In the studied case, such parameter uncertainties drastically reduced the chance of tumor elimination to less than 10%. Additional *in silico* experiments can be conducted to identify, for example, the smallest dose to increase success chance in view of a setting with possible uncertainties. We have also shown that fractionation of dose appears to be as effective as a single dose, and the rest periods between infusions might favor long-term immunological memory. These results corroborate with previous clinical trials using fractioned CAR T cell dose with similar effectiveness to single-dose and persistence of CAR T cells on the blood 20 months after therapy [25].

We identified that uncertainties associated with the tumor proliferation and ability to inhibit the CAR T cells, and CAR T cell proliferation and death are the most significant to therapy success in eliminating the tumors. This opens room for investigating other chimeric antigen T-cell receptors with different target antigen affinities and the blockade of immune checkpoints to boost efficacy and safety. In our model, we did not consider CAR T affinity for each antigen as an explicit parameter, considering it as a result of tumor lysis by CAR T cells. Another aspect that we did not take on consideration is the toxicity effect of CAR T cell therapy (cytokine release syndrome - CRS) because our model is based on an immunodeficient mouse model that lacks this effect. For human data, Hanson et al. [18] made a mathematical model to CAR T cell therapy for B-ALL emphasizing cytokines and CRS, but also considering CAR T effector and memory cells.

Overall, the developed mathematical model may help to shed light on the structure of the treatment protocol. To this end, the model must be calibrated by using one *in vivo* experimental data describing the tumor growth without and with treatment, and *in vitro* lysing data. Once calibrated, the model allows exploring alternative ways of scheduling and infusion dose in view of the current setting specificities, including parameter uncertainties, so as to elicit the one with a higher chance of success. The model provides an *in silico* tool for assessing different issues associated with the therapy such as how CAR T cell dosing can be adjusted according to tumor burden, CAR T cell infusion protocols, immuno-suppressive mechanisms, among others, without further *in vivo* experiments.

## Acknowledgments

The authors thank the support granted by FAPERJ and CAPES.

