## Supplementary Material for "Three-Compartment Model of CAR T-cell Immunotherapy"

### Data used for inferences

We used data from [8] and [5] for parameter inference. We extracted data from these sources using the free software G3Data Graph Analyzer [3] and organized them in the tables shown below.

**Table 1** HDLM-2 tumor burden (data extracted from [8], Figure 4B).

| Day | HDLM-2 [BU] |
|---|---|
| 48 | $9.22 \pm 4.27 \times 10^6$ |
| 55 | $1.09 \pm 0.36 \times 10^7$ |
| 71 | $1.01 \pm 0.40 \times 10^8$ |
| 85 | $2.26 \pm 0.68 \times 10^8$ |
| 92 | $2.02 \times 10^8$ |
| 97 | $9.51 \pm 6.09 \times 10^8$ |
| 108 | $1.31 \times 10^9$ |
| 112 | $2.10 \times 10^9$ |
| 120 | $1.26 \times 10^9$ |
| 136 | $2.73 \times 10^9$ |
| 142 | $7.55 \times 10^9$ |

**Table 2** Cytotoxic activity against HDLM-2 (data extracted from [8], Figure 3B).

| Effector to target ratio | Live cells |
|---|---|
| 0.3 : 1 | $2.46 \times 10^6$ |
| 0.6 : 1 | $9.90 \times 10^5$ |
| 1.25 : 1 | $3.41 \times 10^5$ |
| 2.5 : 1 | $1.82 \times 10^5$ |
| 5 : 1 | $1.62 \times 10^5$ |
| 10 : 1 | $9.65 \times 10^4$ |

**Table 3** Tumor burden (data extracted from [5], Figure 2C).

| Day | Raji-IDO / Raji-control [BU] |
|---|---|
| 0 | $1.05 \pm 0.07 \times 10^8$ |
| **Day** | **Raji-control + CAR T 19 [BU]** |
| 4 | $1.6 \pm 2.1 \times 10^8$ |
| 7 | $5.5 \pm 7.4 \times 10^8$ |

**Table 4** Tumor burden for Raji-IDO treated with CAR T 19 and with CAR T 19+1-MT (data extracted from [5], Figure 3B).

| Day | Raji-IDO + CAR T 19 [BU] |
|---|---|
| 4 | $4.98 \pm 2.01 \times 10^8$ |
| 7 | $16.82 \pm 12.71 \times 10^8$ |
| **Day** | **Raji-IDO + CAR T 19 + 1-MT [BU]** |
| 4 | $1.9 \pm 1.3 \times 10^8$ |
| 7 | $6.9 \pm 4.9 \times 10^8$ |

**Table 5** Cytotoxic activity against Raji-control and Raji-IDO (data extracted from [5], Figure 3D).

| Effector to target ratio | Cytotoxicity [%] |
|---|---|
| 40 : 1 | 73 |
| 20 : 1 | 69 |
| 10 : 1 | 63 |
| 5 : 1 | 56 |

### Inference of the values of model parameters

Parameter estimation is performed for each scenario of interest, from the indirect measurements performed in [8] or in [5]. In this work, we use the Bayesian approach for the parameter inference. The Bayesian approach for estimating the parameters of a model allows us to consider both the uncertainties in the model and the experimental data [6, 9, 4]. A detailed tutorial on this approach focusing on tumor growth models is presented in [2]. The basis of the Bayesian calibration is the Bayes theorem. To present it in the context of the desired estimation, let $\boldsymbol{\theta} = \{\theta_1, \ldots, \theta_k\}$ be the vector of $k$ parameters we want to estimate and $\boldsymbol{d} = \{d_1, \ldots, d_n\}$ the vector of $n$ experimental data. These data may be, for example, the number of tumor cells measured by BLI [1] at $n$ different times. Like every measure, each one is accompanied by uncertainties $\{\sigma_i\}_{i=1}^n$, assumed Gaussian. Let $\boldsymbol{\Sigma}$ be the associated covariance matrix. With these definitions, Bayes' theorem states that:

$$p_{post}(\boldsymbol{\theta}|\mathbf{d}) \propto L(\mathbf{d}|\boldsymbol{\theta}) p_{prior}(\boldsymbol{\theta}) \ , \tag{1}$$

where

- $p_{prior}$ is the prior probability distribution corresponding to our *a priori* knowledge on $\boldsymbol{\theta}$;
- $L$ is the likelihood function; it is a conditional function which allows updating the knowledge about the parameters from the experimental data $\mathbf{d}$;
- $p_{post}$ is the joint posterior probability distribution, that represents the *a posteriori* knowledge about the parameters after we know the data.

In this way, Bayes' theorem updates the *a priori* knowledge, integrating information from data. The result is the joint posterior probability distribution $p_{post}$ which represents the improvement of knowledge about the values of the parameters. The distributions of each parameter are the marginal distributions, wherein the most likely estimates (MLE) are the most probable values.

For the definition of the likelihood function, we assume that the data are independent and that the uncertainties in the measurements follow the Gaussian distribution as mentioned above. Under these conditions, $L$ is given by:

$$L(\mathbf{d}|\boldsymbol{\theta}) = \left((2\pi)^n \ |\boldsymbol{\Sigma}|\right)^{-1/2} \exp\Big( -\frac{1}{2}(\boldsymbol{d}_m - \boldsymbol{d})^t \ \boldsymbol{\Sigma}^{-1}(\boldsymbol{d}_m - \boldsymbol{d})\Big), \tag{2}$$

in which $\boldsymbol{d}_m$ is the vector of the simulated values obtained with the model referring to the experimental measurements using $\boldsymbol{\theta}$. Note that $L(\mathbf{d}|\boldsymbol{\theta})$ provides a measure of the knowledge brought by the data on parameter values for the model. Thus, the joint posterior probability distribution is determined as

$$p_{post}(\boldsymbol{\theta}|\mathbf{d}) \propto \exp\Big(-\frac{1}{2}(\boldsymbol{d}_m - \boldsymbol{d})^t\ \boldsymbol{\Sigma}^{-1}(\boldsymbol{d}_m - \boldsymbol{d})\Big)p_{prior}(\boldsymbol{\theta})\ . \tag{3}$$

In the experiments performed in this work, we assumed equiprobable values in certain ranges for the prior probability distributions because we do not know any prior quantitative knowledge about them. This means that each calibrated parameter follows a uniform *a priori* probability distribution over interval $(a, b)$, denoted by $\mathcal{U}(a, b)$. As the analytical solution of (3) is restricted to particular cases, $p_{post}$ is obtained via Markov Chain Monte Carlo sampling procedures, usually requiring a large number of model simulations. In this work, we used QUESO (*Quantification of Uncertainty for Estimation, Simulation, and Optimization*), a free library for uncertainty quantification [7]. We used a multi-level Monte Carlo method called DRAM, together with the directives available in the QUESO manual and 20,000 samples. Experimental data were extracted by using the software G3Data Graph Analyzer ([3]). Tumor growth parameters ($r$ and $b$) were calibrated considering the part of the proposed model associated with the growth of the tumor in the absence of immunotherapy, i.e., $dT/dt = rT(1 - bT)$. Likewise, the cytotoxicity of the CAR T cells parameter ($\gamma$) was calibrated considering the model $dT/dt = -\gamma C_T T$.

Parameter inference for the HDLM-2 + CAR T 123 scenario

To estimate the parameters $r$ and $b$, we extracted data from Figure 4-B from [8], shown in Table 1. As these data represent untreated tumor growth, we calibrate $r$ and $b$ considering only the tumor growth, i.e., using $\dfrac{dT}{dt} = rT(1-bT)$. The uniform priors were set as: $r \sim \mathcal{U}(0,01\ ,\ 0,1)$ $day^{-1}$, $b \sim \mathcal{U}(10^{-13}, 10^{-10})$ $cell^{-1}$, and $T_0 \sim \mathcal{U}(10^6, 10^8)$ *cell*. The histograms associated with the posterior distributions for $r$ and $b$ are presented in Figure 1(a,b), together with their MLEs. The data used to estimate $\gamma$ came from the standard *in vitro* 4-hour chromium-51 release assay data depicted in Figure 3-B from [8] and shown in Table 2. As described in [8], co-cultures of $2 \times 10^6$ cancer cells and CAR T 123 cells in different proportions were generated. After 4 hours, the number of tumor cells remaining alive was evaluated. To adjust these data, we consider only tumor mortality due to the presence of effector cells, which leads to $\dfrac{dT}{dt} = -\gamma C_T T$. The solution of this equation at 4 hours is used to estimate $\gamma$ using Queso. We used $\gamma \sim \mathcal{U}(10^{-8}, 10^{-5})$ as a prior distribution, which yielded $\gamma_{MLE}$ and the histogram of the posterior distribution shown in Figure 1(c).

Since experimental data from CAR T cells and memory T cells were not available in [8], all the other parameters were estimated through simulations of the three-compartment model. This means that extensive simulations were performed, fixing $r$, $b$, and $\gamma$, until finding a set of parameters that can depict a fast decrease of the tumor after treatment and the tumor elimination after the challenge. The final parameter values used in the simulations are shown in Table 6. Although good representations of the desired scenarios were obtained, it is important to remark that parameter estimation in this way is not unique and may be refined on the lights of more informative data.

Parameter inference for the Raji + CAR T 19 scenario

Raji-like tumors (CD19 + lymphoma) expressing or not the IDO enzyme are quite aggressive. None of the available experiments presented in [5] was able to eliminate the tumor, which shows a clearly exponential growth during the seven days corresponding to the monitoring period.

The data available in [5] are very scarce, referring to only two measurements after treatment, which can hinder or prevent the model calibration. To circumvent this difficulty, we reduce the number of parameters by grouping the rates of proliferation, natural decay, and differentiation of the CAR T cells into a single rate that reproduces the balanced effect of these mechanisms. Moreover, as there is no evidence in the data showing the decrease of the tumor growth rate due to competition for resources, the exponential growth model is used here instead of the logistic model. With these simplifications, we reduced the number of parameters to seven.

All BLI measurements were performed only on the day the treatment is administered and on two subsequent days, providing few data for calibration, which can hinder or prevent the model calibration. To avoid misleading estimation due to lack of data, we simplified the proposed model by grouping parameters on the $C_T$ dynamics and not considering saturation of the tumor growth, yielding

$$\begin{aligned}\frac{dC_T}{dt} &= -\mu C_T + \theta T C_M - \alpha T C_T\\ \frac{dC_M}{dt} &= \beta C_T - \theta T C_M - \mu_M C_M\\ \frac{dT}{dt} &= rT - \gamma T C_T\end{aligned}$$

Thus, in the present representation of the developed model, we set $\mu = \phi - \rho - \mu_T$ and $\beta = \epsilon\rho$, and considered the exponential dynamics of the tumor growth as indicated by the experimental data. The balancing effect of these three mechanisms naturally appears in the estimate of parameter $\mu$.

Like shown in the previous section, $r$ and $\gamma$ were estimated using the untreated tumor growth model $\left(\frac{dT}{dt} = rT\right)$ and the tumor decay model due to effector cells $\left(\frac{dT}{dt} = -\gamma C_T T\right)$, respectively. The uniform priors were set as:

$r \sim \mathcal{U}(0,1\ ,\ 0,9)\ day^{-1}$, and $\gamma \sim \mathcal{U}(10^{-9}, 10^{-7})\ (cel \cdot day)^{-1}$. The histograms associated with the posterior distributions for $r$ and $\gamma$ are presented in Figure 2(a,b), together with their MLEs. Since there is no data for CAR T cells (effector or memory), we estimated all the reminder parameters through extensive simulations for the Raji-control tumor to represent the behavior of the tumor growth depicted in Figure 2C from [5] (Table 3). Once this set of parameters were obtained, we fixed all parameters but $\alpha$. All the obtained parameters are shown in Table 6. The $\alpha$ parameter, associated with the mechanism by which tumor cells inhibit effector cells, was calibrated using QUESO and in all the following situations we used the prior $\alpha \sim \mathcal{U}(10^{-8}, 3 \times 10^{-8})(cell \cdot day)^{-1}$. For Raji-control + CAR T 19, using the data shown in Table 3, the histogram associated with the posterior distribution for $\alpha$ is presented in Figure 2(c), together with its MLE that amounts $\alpha_{MLE} = 1.25 \times 10^{-8}(cell \cdot day)^{-1}$.

For the Raji-IDO tumor, inference of parameter $\alpha$ is performed for the cases with and without 1-MT, using the experimental data presented in Figure 3B from [5] (Table 4). As mentioned before, in these inferences only the parameter $\alpha$ is calibrated to capture the effect of the immunosuppression effect, keeping all other parameter values as set in Table 6. The MLEs obtained for the treatments with Raji-IDO + CAR T 19 and Raji-IDO + CAR T 19 + 1-MT are $\alpha_{MLE} = 1.46 \times 10^{-8}\ (cell \cdot day)^{-1}$ and $\alpha_{MLE} = 1.26 \times 10^{-8}\ (cell \cdot day)^{-1}$, respectively. Note the decrease of $\alpha_{MLE}$ when the IDO inhibitor is used and the similaty between the $\alpha_{MLE}$ values obtained for Raji-control + CAR T 19 and Raji-IDO + CAR T 19 + 1-MT.

### Supplementary Table and Figures

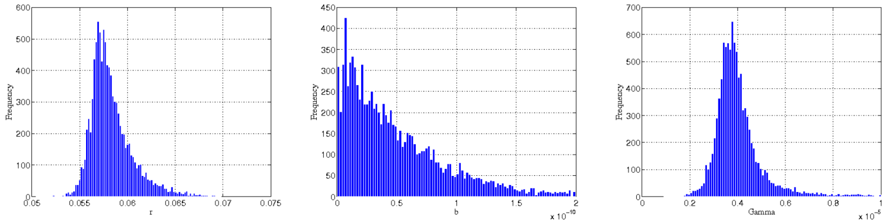


(a) $r_{MLE} = 5.65 \times 10^{-2} day^{-1}$ (b) $b_{MLE} = 1.40 \times 10^{-12} cell^{-1}$ (c) $\gamma_{MLE} = 3.72 \times 10^{-6} (cell \cdot day)^{-1}$

**Fig. 1** HDLM-2 + CAR T 123 scenario: posterior histograms of $r$ (a), $b$ (b), and $\gamma$ (c). The maximum *a posteriori* estimates of marginal distributions are indicated by the subscript MLE. Since all priors are uniform distributions, there was a substantial improvement of the knowledge about the parameters with the application of Bayesian inference, with parameters presenting log-normal like posterior distributions. However, the inference of the inverse of the carrying capacity ($b$) presents greater uncertainty which may be explained by the lack of saturation of the tumor burden for the untreated mice (see Figure 4B of [8]).

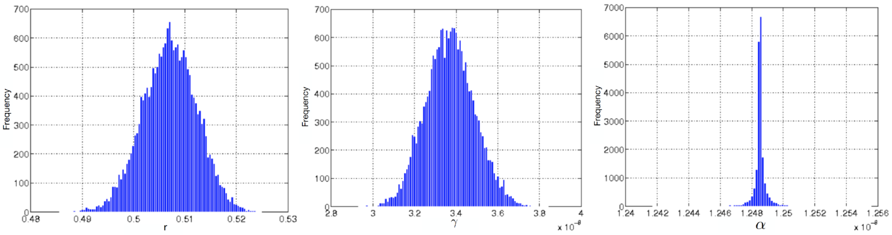


(a) $r_{MLE} = 5.07 \times 10^{-1} day^{-1}$ (b) $\gamma_{MLE} = 3.37 \times 10^{-8} (cell \cdot day)^{-1}$ (c) $\alpha_{MLE} = 1.25 \times 10^{-8} (cell \cdot day)^{-1}$

**Fig. 2** Raji-control + CAR T 19 scenario: posterior histograms of $r$ (a), $\gamma$ (b), and $\alpha$ (c). The maximum *a posteriori* estimates of marginal distributions are indicated by the subscript MLE. As in the case of HDLM-2 cancer, there was a substantial improvement of the knowledge about the parameters with the application of Bayesian inference, with the calibrated parameters also presenting log-normal like posterior distributions.

**Table 6** Model parameter values. The parameters estimated using QUESO are indicated with $\star$, whose values are the MLEs.

| **HDLM-2 + CAR T 123** | | **Raji-control + CAR T 19** | |
|---|---|---|---|
| **Parameter** | **Value** | **Parameter** | **Value** |
| $\phi$ | $0.265\ day^{-1}$ | | |
| $\rho$ | $5.0 \times 10^{-2}\ day^{-1}$ | $\mu$ | $5.36 \times 10^{-5}\ day^{-1}$ |
| $\mu_T$ | $3.0 \times 10^{-1}\ day^{-1}$ | | |
| $\epsilon$ | 3.0 | $\beta$ | $1.6\ day^{-1}$ |
| $\theta$ | $6.0 \times 10^{-6}\ (cell \cdot day)^{-1}$ | $\theta$ | $2.3 \times 10^{-4}\ (cell \cdot day)^{-1}$ |
| $\alpha$ | $4.5 \times 10^{-8}\ (cell \cdot day)^{-1}$ | $\alpha^\star$ | $1.25 \times 10^{-8}\ (cell \cdot day)^{-1}$ |
| $\mu_M$ | $5.0 \times 10^{-3}\ day^{-1}$ | $\mu_M$ | $6.89 \times 10^{-7}\ day^{-1}$ |
| $r^\star$ | $5.65 \times 10^{-2}\ day^{-1}$ | | |
| $b^\star$ | $1.40 \times 10^{-12}\ cell^{-1}$ | $r^\star$ | $5.07 \times 10^{-1}\ day^{-1}$ |
| $\gamma^\star$ | $3.72 \times 10^{-6}\ (cell \cdot day)^{-1}$ | $\gamma^\star$ | $3.37 \times 10^{-8}\ (cell \cdot day)^{-1}$ |
